# Classical metapopulation dynamics and eco-evolutionary feedbacks in dendritic networks

**DOI:** 10.1101/033639

**Authors:** Emanuel A. Fronhofer, Florian Altermatt

## Abstract

Eco-evolutionary dynamics are now recognized to be highly relevant for population and community dynamics. However, the impact of evolutionary dynamics on spatial patterns, such as the occurrence of classical metapopulation dynamics, is less well appreciated. Here, we analyse the evolutionary consequences of spatial network connectivity and topology for dispersal strategies and quantify the eco-evolutionary feedback in terms of altered classical metapopulation dynamics. We find that network properties, such as topology and connectivity, lead to predictable spatio-temporal correlations in fitness expectations. These spatio-temporally stable fitness patterns heavily impact evolutionarily stable dispersal strategies and lead to eco-evolutionary feedbacks on landscape level metrics, such as the number of occupied patches, the number of extinctions and recolonizations as well as metapopulation extinction risk and genetic structure. Our model predicts that classical metapopulation dynamics are more likely to occur in dendritic networks, and especially in riverine systems, compared to other types of landscape configurations. As it remains debated whether classical metapopulation dynamics are likely to occur in nature at all, our work provides an important conceptual advance for understanding the occurrence of classical metapopulation dynamics which has implications for conservation and management of spatially structured populations.

## Introduction

Evolution is recognized to be rapid enough to affect ecological dynamics, which may lead to eco-evolutionary feedbacks (Hanski, 2012; Ellner, 2013). Although a majority of species on earth live in fragmented habitats and therefore form spatially structured populations, most of eco-evolutionary research has focused on single, isolated populations and communities in a non-spatial context (for a recent review, see Koch et al., 2014). It remains therefore less well appreciated that evolutionary dynamics affect classical spatial patterns, such as the dynamics of populations living in networks of interconnected local patches, that is, metapopulations and metacommunities.

The classical metapopulation concept (Levins, 1969; Hanski and Gaggiotti, 2004), and the notion that most natural populations are spatially structured, has extensively influenced decades of research in spatial ecology and conservation (e.g., Driscoll et al., 2014). More recently, spatial structure has found its way into community (Leibold et al., 2004) and ecosystem research (Loreau et al., 2003). While the metapopulation, –community and –ecosystem concepts are at the heart of spatial ecology, most theoretical and conceptual work on these spatial systems has still not included an explicit description of space (but see Hanski, 2001; Hanski et al., 2004; Baguette et al., 2013). Space, and more specifically spatial network configuration and inter-patch connectivity, is often included only implicitly, for example, by assuming global dispersal abilities (e.g., Poethke et al., 2011; Weigang and Kisdi, 2015) or only considering two patches (e.g., McPeek and Holt, 1992; Amarasekare, 2004). Even when space is considered explicitly, often simplistic connectivity patterns are assumed, such as grid-based, nearest-neighbour dispersal (e.g., Travis and Dytham, 1998; Kubisch et al., 2015). However, these assumptions are most likely erroneous in any natural spatially structured population, community or ecosystem.

Existing research on consequences of alternative network connectivities and topologies suggests that these properties are of pivotal importance for ecological and evolutionary dynamics. For example, (Bascompte and Sole, 1996) Fagan (2002), Vuilleumier and Possingham (2006), Labonne et al. (2008), Gilarranz and Bascompte (2012) and Shtilerman and Stone (2015) have studied the effects of network topology, respectively symmetry on metapopulation viability and persistence. They found that network structure impacts demography and leads to higher extinction probabilities than otherwise expected. In a multi-species context, Carrara et al. (2012) and Seymour et al. (2015) have demonstrated that spatial and temporal patterns of biodiversity are impacted by the specific connectivity pattern of a landscape (see also Holland and Hastings, 2008; Salomon et al., 2010). Generally, these findings suggest that branching networks may support higher levels of biodiversity in comparison to more simply structured landscapes. In analogy, Morrissey and de Kerckhove (2009) and Paz-Vinas and Blanchet (2015) have shown that network topology heavily impacts genetic diversity. Recently, (Muneepeerakul et al., 2011) and Henriques-Silva et al. (2015) have reported that network topology may even impact the evolution of dispersal kernels respectively density-dependent dispersal strategies in metapopulations. While these studies have addressed important aspects of spatial ecology and evolution, no study to date has integrated the individual elements, to how these eco-evolutionary dynamics and feedbacks impact the likelihood over observing classical metapopulation dynamics.

Using a coherent eco-evolutionary framework, we investigate theoretically how evolutionary and ecological dynamics interact in networks of populations with different connectivity and topology. We focus on the evolution of dispersal, as this trait has been shown to be evolving in a wide range of taxa (e.g., Phillips et al., 2006; Saastamoinen, 2008; Fronhofer et al., 2014; Fronhofer and Altermatt, 2015), and to centrally influence the dynamics of spatially structured populations. Specifically, we ask how the evolution of dispersal in networks of varying connectivity and topology impacts the occurrence of classical metapopulation dynamics.

Our interest in exploring the occurrence of classical metapopulation dynamics stems from the current debate on whether these dynamics occur at all in natural systems (among others, Baguette, 2004; Driscoll, 2007, 2008; Driscoll et al., 2010). A range of alternative scenarios, including mainland-island, source-sink or panmictic spatially structured systems seem to be possible (Harrison, 1991), but would all lead to altered system properties such as extinction probabilities, number of occupied patches (occupancy), number of extinctions and recolonizations (turnover), and genetic structure (the fixation index, *F_ST_*). Clearly, appropriate conservation and management strategies must take these differences into account.

While our theoretical considerations are, in principle, valid for any type of terrestrial or aquatic network of patches, we apply our findings to a classical example of habitat networks: dendritic, riverine systems. Rivers are not only very diverse and of high significance with respect to ecosystem services (Vörösmarty et al., 2010), but they also have an inherent dendritic network structure which drives dispersal and diversity patterns (Muneepeerakul et al., 2008; Grant et al., 2007; Altermatt, 2013; Mari et al., 2014). Furthermore, riverine ecosystems are an especially interesting testbed for theoretical predictions regarding the consequences of network properties, as they are currently experiencing large changes in network configuration and connectivity by ongoing fragmentation, dam and channel building (Grill et al., 2015; Grant et al., 2012).

We find that network topology and connectivity lead to predictable, spatio-temporally correlated, patterns of fitness expectations, which alter evolutionarily stable (ES) dispersal strategies and lead to eco-evolutionary feedbacks on landscape level metrics. Dendritic networks, and especially riverine connectivity patterns, thereby favour the emergence of classical metapopulation dynamics. In comparison to such dendritic spatial structures, classical metapopulation dynamics are less likely found in symmetric networks, which are often assumed in metapopulation models. In the context of the ongoing debate regarding the occurrence of classical metapopulation dynamics in natural systems (among others, Baguette, 2004; Driscoll, 2007, 2008; Driscoll et al., 2010), our findings highlight the significance of network connectivity and topology for spatial ecological and evolutionary dynamics.

## Model description

### General overview

We use a general, stochastic simulation model of a spatially structured population of individuals living in distinct habitat patches with local competition for resources and non-overlapping generations (see, e.g., Fronhofer et al., 2013, 2014). Local populations are connected by dispersal, which is defined by every individual’s dispersal rate and by the landscape’s topology, that is, the spatial arrangement of habitat patches (connectivity matrix, see, e.g., Seymour et al., 2015). Dispersal is natal. Importantly, the dispersal trait is heritable and subject to evolution.

Using network topologies that either only differ in connectivity (i.e. number of links from one patch to other patches) or in topology (regular, grid-like networks versus branching, dendritic networks), we explore the eco-evolutionary consequences of network structure on dispersal evolution and metapopulation dynamics, measured as occupancy (*O*, the relative number of occupied patches), turnover (*T*, the relative number of extinctions and recolonizations) and genetic structure (*F_ST_*, which captures variation in allele frequencies among populations). Following Hanski et al. (1995) and Fronhofer et al. (2012), we define classical metapopulations as any spatially structured population that shows less than 90% occupancy, more than 5% turnover and a global *F_ST_* ≥ 0.1. To relate our results to real-world systems, we supplement our general analysis with the example of riverine networks, which typically exhibiting dendritic network structure, also including characteristic variation in habitat size (carrying capacity; Rodriguez-Iturbe and Rinaldo, 1997) and downstream water flow.

### Landscape

We assume a lattice-type spatial network, in which nodes are habitat patches and links are uninhabited “matrix”, reflecting a spatially structured population in a network. For an overview of continuous space networks see Grant et al. (2007). We analyse the eco-evolutionary consequences of three types of landscapes, that all have 36 nodes (patches) for comparability:

1. Lattice landscapes with varying connectivity, that is, links per node (see Fig. 1 top row for a graphical representation). Our choice covers the two extreme possibilities, namely a fully connected network (often termed “global dispersal”) where every node connects to every other node (maximal number of links), and a circular network where every node has only two links. We additionally explore two intermediate cases: a network allowing for nearest neighbour dispersal to the eight nearest neighbours (NN8, Moore neighbourhood) and one allowing for dispersal to the four nearest neighbours (NN4, von Neumann neighbourhood) on a regular grid.
2. We further analyse the effect of topology by comparing the dynamics in the lattice landscapes with a number of bifurcating networks (see Fig. 4 top row, two networks on the left, for a graphical representation): we use one landscape which is dendritically bifuracting in analogy to Morrissey and de Kerckhove (2009) and complement our analysis with five realizations of fractal dendritic landscapes (OCNs; optimal channel networks; we here use the same OCNs as Carrara et al., 2014).
3. We finally use an OCN in combination with the characteristic riverine distribution of carrying capacities found in nature (i.e., carrying capacities increasing form up-to downstream patches; Rodriguez-Iturbe and Rinaldo, 1997; Carrara et al., 2014) and two strengths of downstream flow (probability of up-/downstream dispersal: 0.5/0.5 and 0.1/0.9) in order to explore the robustness of our findings in more realistic settings and for the most characteristic example of dendritic networks, namely rivers (see Fig. 4 top row, two networks on the right, for a graphical representation). The typically riverine distribution of carrying capacities in nature results from the drainage area of the respective patches as described by Carrara et al. (2014). Based on their drainage area, patches were assigned to four relative size categories (1, 1.75, 3 and 6). In order to keep the simulations comparable, we kept the landscape-level carrying capacity (sum of all local carrying capacities) for all landscapes constant, while assigning local carrying capacities according to their relative sizes. For a more detailed description refer to Carrara et al. (2014).

**Figure 1:**
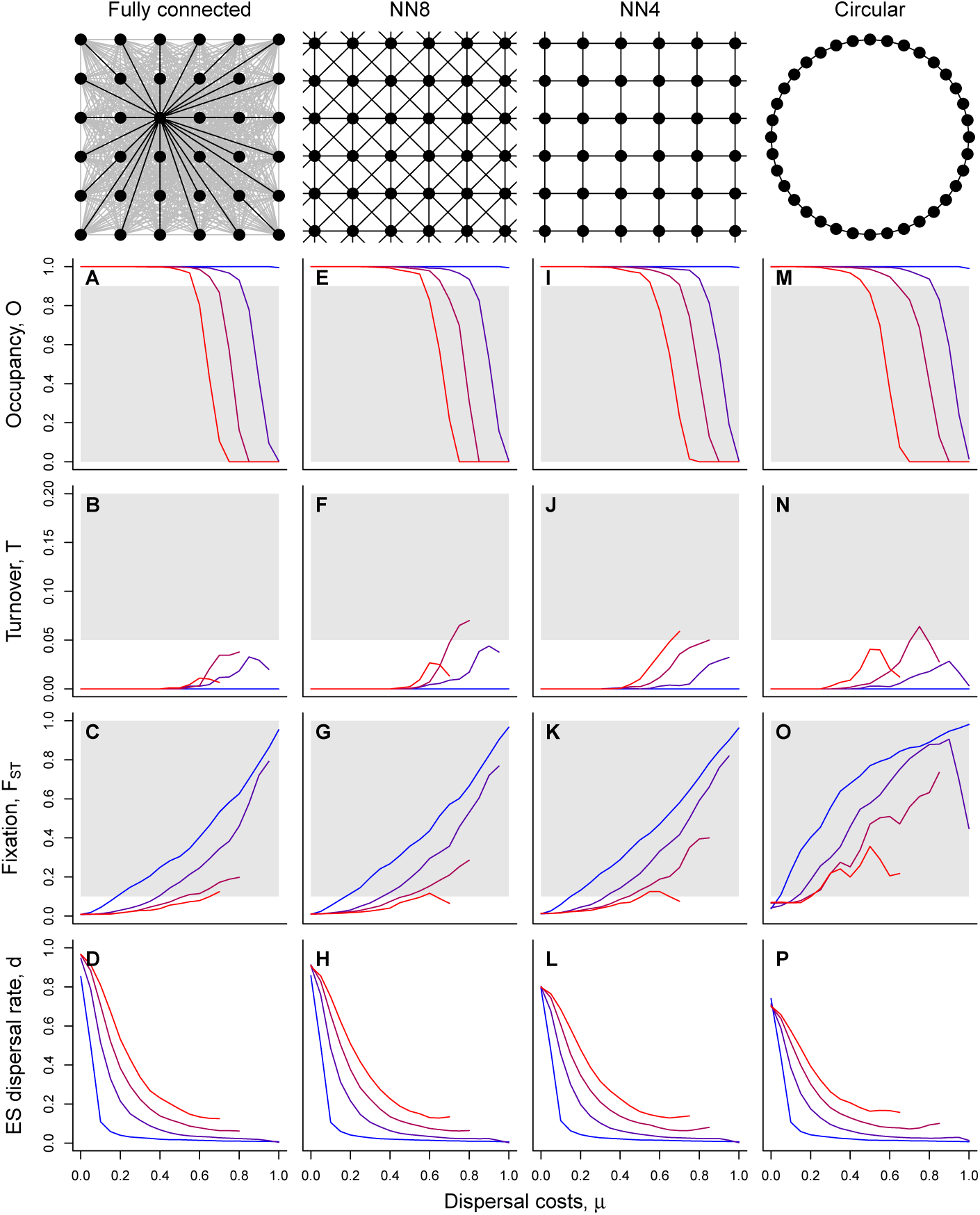
Ecological and evolutionary dynamics of spatially structured populations with different degrees of connectivity. Environmental stochasticity (*σ*) increases from blue to red (*σ* ∈ {0, 1, 1.5, 2}). Grey areas indicate values of occupancy (*O*), turnover (*T*) and genetic fixation (*F_ST_*) typically assumed to be characteristic of classical metapopulations (Fronhofer et al., 2012), that is, *O* ≤ 0.9, *T* ≥ 0.05 and a global *F_ST_* ≥ 0.1. Note that, as the curves are relatively steep, the exact choice of values does not critically alter the qualitative results. Fixed parameters: *K* = 100, *σ* ∈{0, 1, 1.5, 2}, *ϵ* = 0, *λ*_0_ = 2. The lines were smoothed with a running mean window of 2. Note that the left network representation in the top row (“Fully connected”) only highlights the connection from one patch to all others; all other connections are analogous.

### Local patch dynamics

Patches can be inhabited by diploid male and female individuals. All individuals are characterized by their dispersal rate (for a detailed description of the dispersal process see below) and by 10 neutral marker loci with 100 alleles, to track population genetic summary statistics such as *F_ST_*. Females mate randomly in their local patch and produce diploid offspring that inherit maternal and paternal dispersal and neutral alleles. During inheritance, traits may change due to mutations. For the dispersal rate this is captured by altering the parental allele value by a random number drawn from a normal distribution with mean zero and standard deviation Δ*m* =0.2 in case of a mutation (mutation rate: *m_dispersal_* = 0.0001; no boundary conditions). For the neutral loci, the mutation rate is *m_neutral_* = 0.00001 and in case of a mutation any of the 100 alleles can be drawn. This allows evolutionary dynamics to occur during the simulations.

We assume local density regulation in all networks, that is, competition acts at the local population level. As our model assumes discrete, non-overlapping generations, we use the logistic growth model provided by Beverton and Holt (1957):

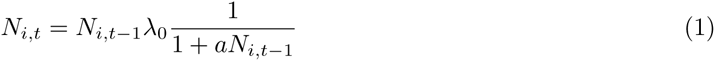

with *i* as the patch number, *t* as the time step and 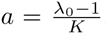. *K* is the local carrying capacity and *λ*_0_ represents the growth rate. Consequently, every female produces a mean of 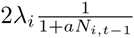 offspring (the multiplication with 2 allows to interpret *λ_i_* as a per capita rate even though males do not reproduce) with a sex ratio of 0.5. The realized number of offspring is drawn from a Poisson distribution in order to capture demographic stochasticity. Our model includes spatio-temporally uncorrelated environmental stochasticity caused by variation in offspring number: for every patch and generation *λ_i_* is drawn from a log-normal distribution with mean *λ*_0_ and standard deviation *σ*.

Finally, we assume that every patch may go extinct at a certain rate (*ϵ*) due to external, density-independent factors, such as catastrophic floods or geologic events. Using non-overlapping generations, all adults die after reproduction and the juveniles start a new generation in the next time step.

### Dispersal

Local patches are linked by dispersal events. We assume dispersal to be natal, that is, to occur before reproduction. Dispersal is defined by the individual dispersal rate and by the landscape’s topology (connectivity matrix). The phenotypic dispersal rate is determined as the mean of an individual’s two dispersal alleles. As these alleles may mutate and we do not assume any boundary conditions, values may be below 0 or above 1. Dispersal phenotypes < 0 and > 1 are rounded to 0 and 1, respectively. This procedure avoids assuming boundary conditions for mutations and the associated biases.

Emigration must not necessarily lead to successful immigration, as we assume dispersal costs (*µ*) in form of dispersal mortality. This mortality term summarize all possible costs related to dispersal, such as time, opportunity, risk or energetic costs (Bonte et al., 2012).

### Numerical analyses

All simulations, with 25 replicates each, were run for 5,000 generations, which allowed the system to reach quasi-equilibrium. The simulations were initialized with fully occupied patches and a sex ratio of 0.5. At initialization, dispersal alleles were randomly drawn between 0 and 1, and neutral alleles were randomly assigned one of 100 possible alleles (integer numbers).

Turnover (*T*) was quantified as the relative number of extinctions and recolonizations after dispersal, which accounts for rescue effects. Occupancy (*O*) is the relative number of occupied patches. Population genetics analyses were performed on the individuals of the last generation (*t* = 5000) with the statistical software package R (version 3.2.0; package “hierfstat” version 0.04–14). The reported values of turnover, occupancy, *F_ST_* and dispersal rates were always measured in the last generation of the simulations and are means over the 25 replicates. See Table 1 for the explored parameter space and the Appendix for a sensitivity analysis.

**Table 1:** Important model parameters, their meaning and tested values. Standard values are underlined.

| Parameter | Values | Meaning |
| --- | --- | --- |
| $K$ | 50, <u>100</u>, 200 | carrying capacity |
| $\sigma$ | 0, 0.25, 0.5, ... , 4 | environmental stochasticity |
| $\epsilon$ | <u>0</u>, 0.05, 0.1 | local patch extinction probability |
| $\lambda_0$ | 1.5, <u>2</u>, 4 | fecundity |
| $\mu$ | 0, 0.05, 0.1, ... , 1 | dispersal costs |

## Results

### Consequences of the degree of connectivity

We found substantial evolutionary effects of network connectivity on the evolutionarily stable (ES) dispersal rate (Fig. 1 D, H, L and P). Landscapes with less connectivity, for example the circular landscape, lead to the evolution of lower dispersal rates. This affects ecological patterns, implying a decrease in occupancy (Fig. 1 A, E, I and M), and an increase in turnover (Fig. 1 B, F J and N). When looking at global *F_ST_* values, we also find consequences for population genetic patterns where the functional relationship between *F_ST_* and dispersal costs (*µ*) changes from convex to concave (Fig. 1 C, G, K and O).

The evolutionary effect of network connectivity on dispersal is explained by an altered spatial kin structure (Fig. 2). A fully connected network does, by definition, not show any spatial kin structure, except that kin competition is usually stronger in the natal patch than elsewhere. We use pairwise *F_ST_* values between patches of origin and potential target patches, which are inversely proportional to relatedness, to illustrate this effect: in a fully connected network the relevant pairwise *F_ST_* value relative to the global *F_ST_* value is, unsurprisingly, 1, which implies no difference between global mean relatedness and relatedness between any pair of origin and target patches. In a spatially explicit network, such as nearest neighbour or circular networks, the relative *F_ST_* is consistently lower, indicating that relatedness between natal and target patches is, on average, increased. Consequently, increased relatedness between origin and target patch populations reduces the advantages of dispersal associated with reducing kin competition (Hamilton and May, 1977) and leads to lower ES dispersal rates.

**Figure 2:**
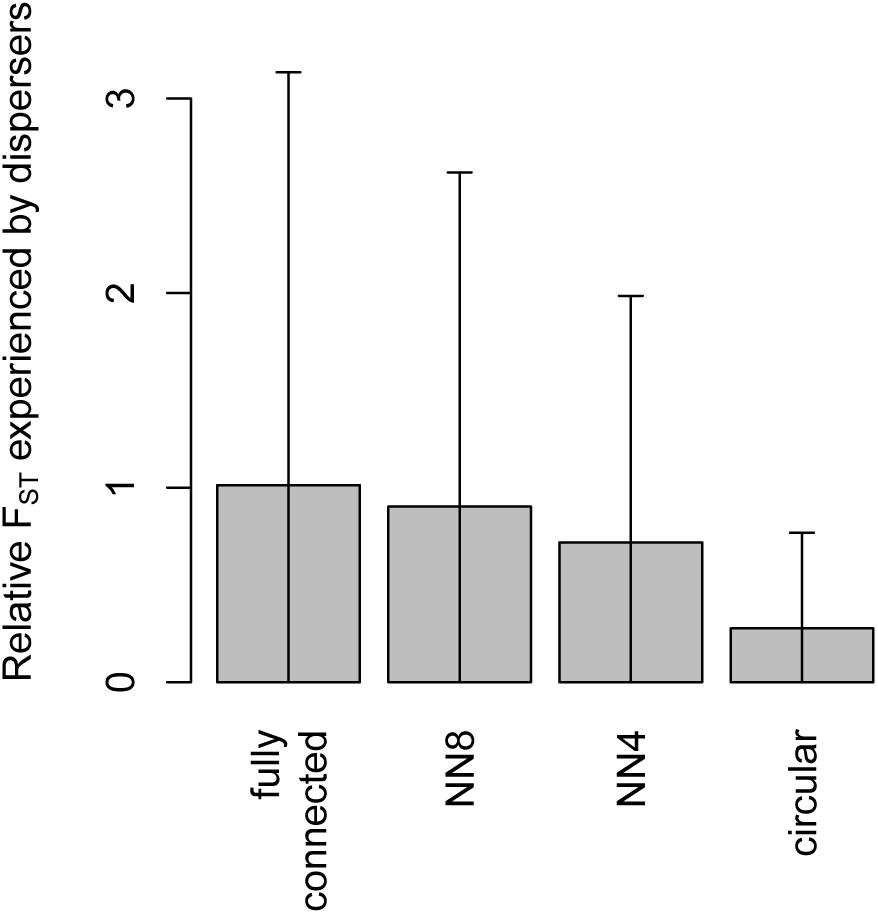
Relative *F_ST_* values experienced by dispersing individuals. To visualize how network connectivity impacts spatial kin structure, we use pairwise *F_ST_* values between origin and target patches (genetic differentiation) relative to the global *F_ST_* value. Note that genetic differentiation is inversely proportional to relatedness. In a fully connected network, without any spatial kin structure, the relevant pairwise *F_ST_* value relative to the global *F_ST_* value is 1. Values < 1 indicate increased kin structure in potential target patches relative to the global average. We show mean ± s.d. values measured at the end of the simulations. Fixed parameters: *K* = 100, *σ* = 0, *ϵ* = 0, *λ*_0_ = 2, *µ* = 0.

The evolutionary effect of network connectivity on dispersal, and the feedback on occupancy, turnover and genetic structure (Fig. 1), has important consequences for the occurrence of classical metapopulation dynamics (Fig. 3). We typically find that increasing dispersal costs (*µ*) and decreasing connectivity (Fig. 1 from A to D) leads to a higher probability of metapopulation extinction due to reduced rescue effects. Metapopulation extinction also increases with increasing environmental stochasticity (*σ*) due to increased local extinctions. Occupancy typically decreases abruptly from 1 (fully occupied) to zero (extinct; see also Fig. 1 A, E, I and M). Therefore, we consistently find a narrow band of intermediate occupancies, which are characteristic for classical metapopulation dynamics, right before the extinction region.

**Figure 3:**
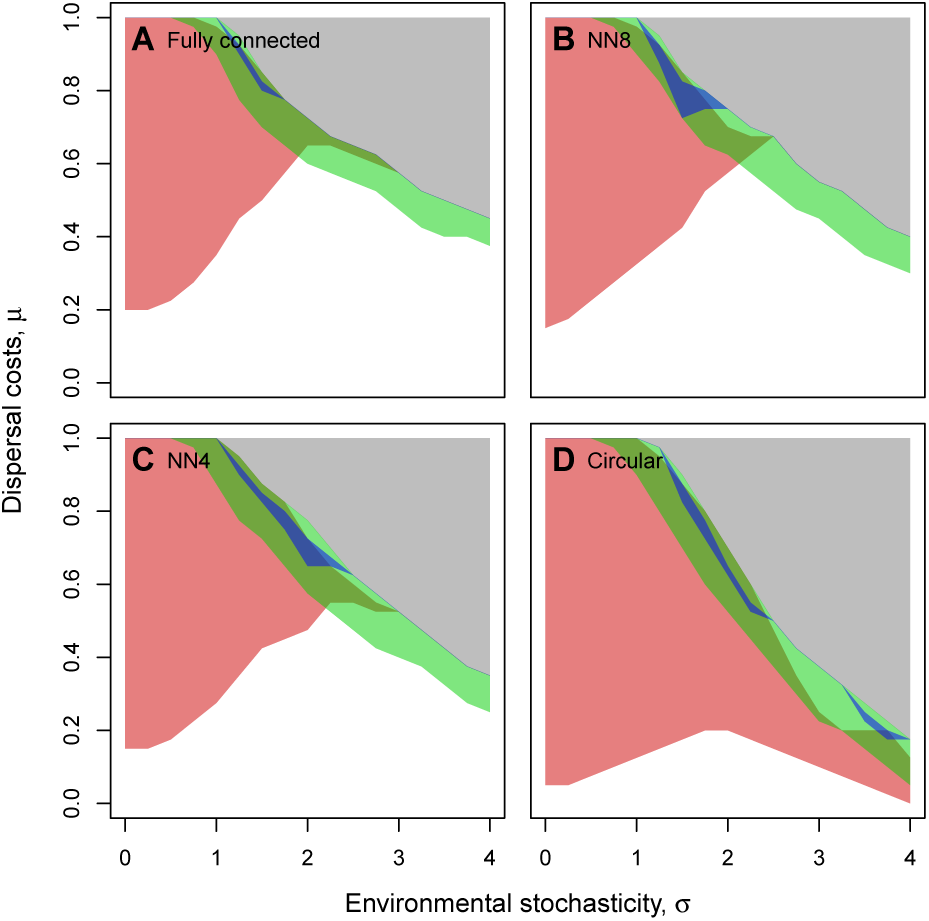
Classical metapopulation dynamics in systems with different degrees of connectivity (for visualisations of the networks, see Fig. 1). Grey: extinction; Red: *F_ST_* ≥ 0.1; Blue: *T* ≥ 0.05; Green: *O* ≤ 0.9. Fixed parameters: *K* = 100, *ϵ* = 0, *λ*_0_ = 2. The polygon lines were smoothed with a running mean window of 2. The original simulation results are reported in the Appendix (Figs. A1 – A4).

Genetic structure (*F_ST_*) typically increases with deceasing ES dispersal rates (Fig. 1 C, G, K and O), that is, with increasing dispersal costs (*µ*) and decreasing environmental stochasticity (*σ*). As decreasing connectivity leads to the evolution of lower dispersal rates, *F_ST_* also increases with decreasing connectivity; more specifically, the shape of the relationship between *F_ST_* and dispersal costs (*µ*) changes from convex to concave (Fig. 1 C, G, K and O). Only in networks with very low connectivities (here: circular; Fig. 3 D) very high values of environmental stochasticity lead to an additional increase in *F_ST_*. This increase in *F_ST_* is due to an increase in local extinctions, leading to founder effects and locally increased drift due to small population sizes. As a result, populations become genetically more differentiated at a global scale.

In general, significant turnover only occurs within the band of intermediate occupancy. However, high environmental stochasticity decreases turnover because such stochasticity selects for increased dispersal which leads to rescue effects. This changes for circular networks (Fig. 1 D): an additional region with increased turnover appears due to the same reasons as *F_ST_* increases.

### Consequences of dendritic topology

Changing network topology from equally connected to bifurcating and dendritic has a similar effect as reducing network connectivity (Fig. 4). However, dendritic networks select even stronger against dispersal than reduced connectivity (Fig. 4 D and H), which, as outlined above, reduces occupancy (Fig. 4 A and E), and increases turnover rates (Fig. 4 B and F) and *F_ST_* (Fig. 4 C and G).

**Figure 4:**
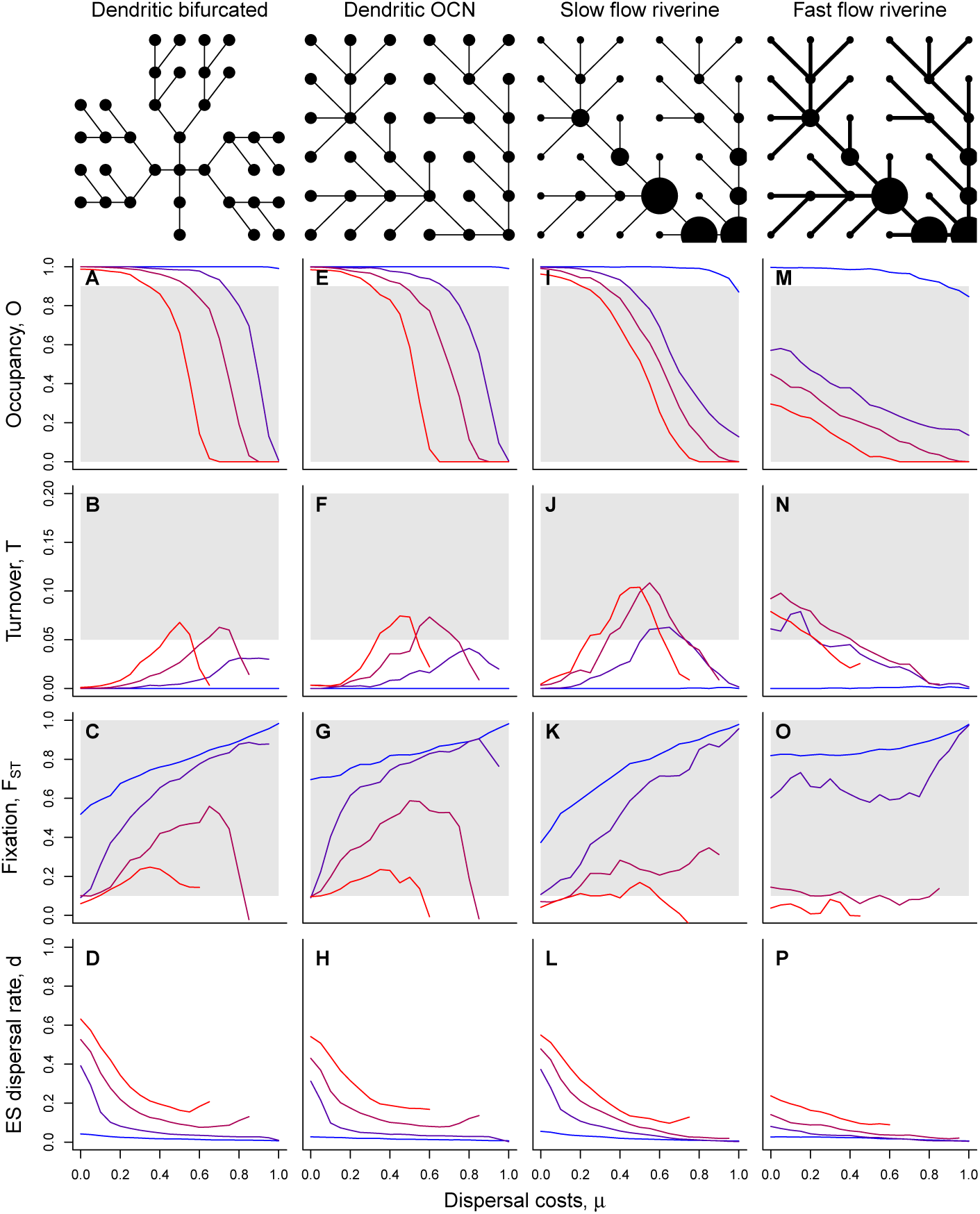
Ecological and evolutionary dynamics of spatially structured populations with different network topologies, including typical riverine networks with different degrees of flow (slow: 0.5 probability of up-or downstream dispersal; fast: 0.1 and 0.9 probability of up-respectively downstream dispersal). Environmental stochasticity (*σ*) increases from blue to red (*σ* ∈{0, 1, 1.5, 2}). Grey areas indicate values of occupancy (*O*), turnover (*T*) and genetic fixation (*F_ST_*) typically assumed to be characteristic of classical metapopulations. Fixed parameters: *K* = 100 (riverine: *K* ∈{57, 99, 170, 340}), *σ* ∈{0, 1, 1.5, 2}, *ϵ* = 0, *λ*_0_ = 2. The lines were smoothed with a running mean window of 2.

While selection for reduced dispersal emerges in systems with low connectivity due to a strong local kin structure (Fig. 2), dendritic networks select for even less dispersal due to an emergent spatial heterogeneity in population densities (Fig. 5). Patches that are less connected typically show lower densities in comparison to well-connected patches. Trivially, this is a result of altered dispersal patterns: patches with only one link usually connect to patches with two or more links, which implies that the earlier patches loose all of their emigrants while they only receive a fraction (based on the number of connections) of immigrants of their neighbouring patch. As a consequence, patches with only one connection have reduced densities as they have more emigrants than immigrants. The opposite is true for the receiving patch. Taken together, the reduced connectivity and the topology of dendritic networks leads, via an eco-evolutionary feedback loop, to the emergence of a larger area of classical metapopulation dynamics (Fig. 6 A and B).

**Figure 5:**
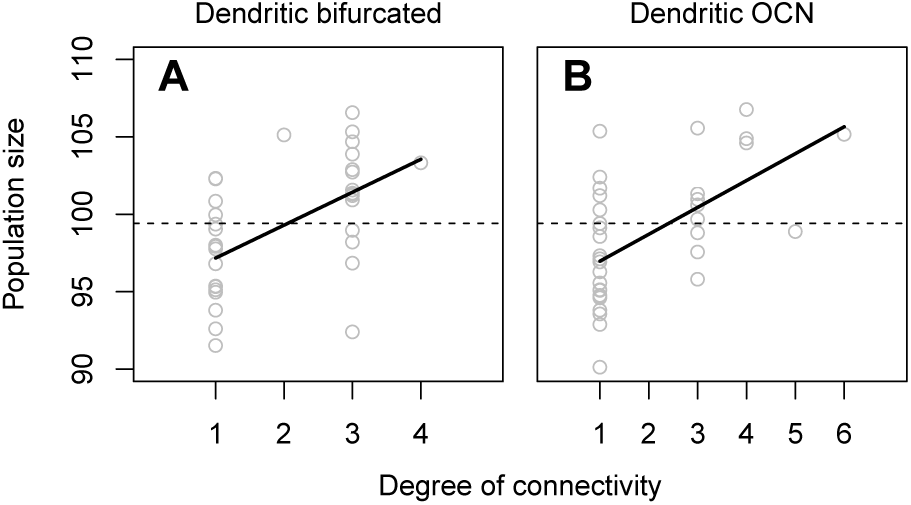
Spatial distribution of population sizes in dendritic networks as a function of patch connectivity. The solid line is a mean-squared regression. The dashed line shows mean population size in a fully connected network. Fixed parameters: *K* = 100, *σ* = 0, *ϵ* = 0, *λ*_0_ = 2, *µ* = 0.

**Figure 6:**
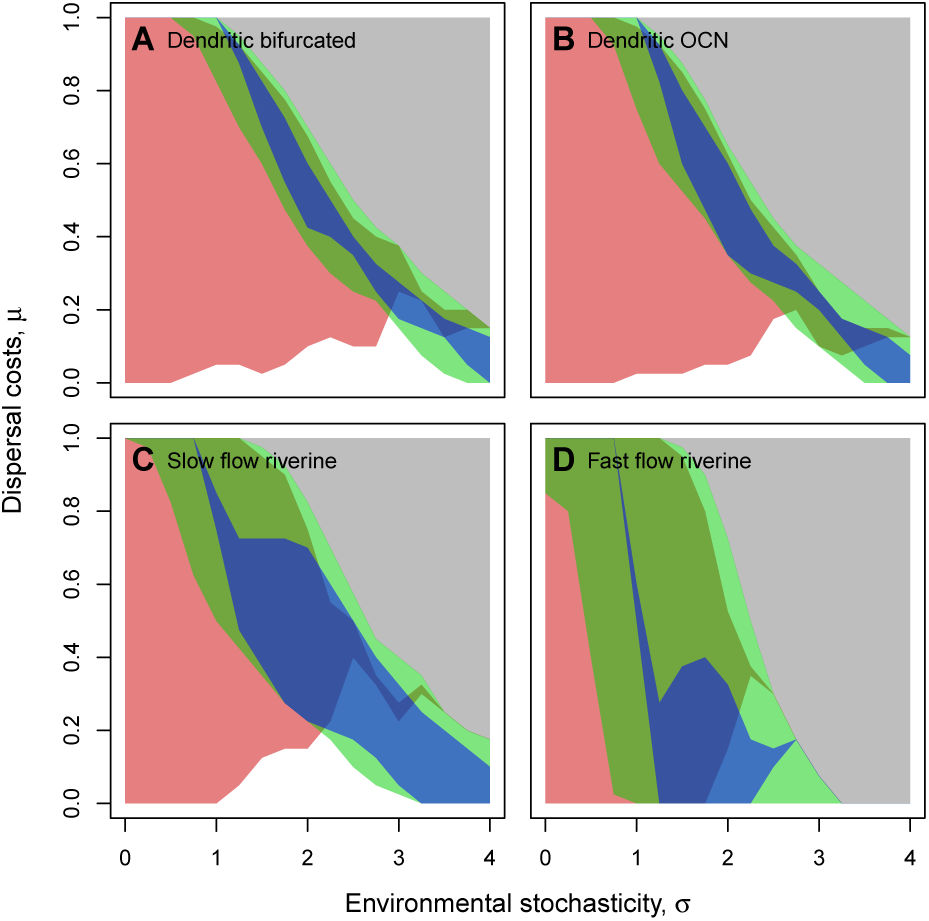
Classical metapopulation dynamics in systems with different network topologies (for visualisations of the networks, see Fig. 4). Grey: extinction; Red: *F_ST_* ≥ 0.1; Blue: *T* ≥ 0.05; Green: *O* ≤ 0.9. Fixed parameters: *K* = 100 (riverine: *K* ∈{57, 99, 170, 340}), *ϵ* = 0, *λ*_0_ = 2. The polygon lines were smoothed with a running mean window of 2. The original simulation results are reported in the Appendix (Figs. A5 and A6).

### Consequences of riverine characteristics

Riverine dendritic networks, characterized by unidirectional flow and a hierarchical distribution of carrying capacities, select even more for low dispersal rates, which generally strengthens all patterns discussed above (Fig. 4 I–P). Consequently, classical metapopulation dynamics emerge across a larger part of parameter space (Fig. 6 C, D).

## Discussion

Our results confirm that the specific network structure, underlying a spatially structured population, has strong eco-evolutionary consequences. Connectivity and topology impact large-scale spatial dynamics and the genetic structure of metapopulations by affecting the evolution of dispersal strategies. Importantly, we show that network structure influences spatial eco-evolutionary dynamics in predictable ways: decreasing connectivity and increasingly dendritic topologies select against dispersal and consequently decrease occupancy, increase turnover and increase the risk of metapopulation extinction.

We find that network topologies with realistic natural analogues, such as rivers, are more likely to exhibit classical metapopulation dynamics than the commonly assumed lattice-like networks (see also Fronhofer et al., 2012). Our findings have direct conservation relevant implications: we suggest that conservation strategies need to better, and system-specifically, incorporate effects of habitat network topology and connectivity, and changes thereof, for the long-term protection of populations and communities. As species in riverine networks exhibit an increased occurrence of classical metapopulation dynamics, they may also be more sensitive to changes in patch availability and connectivity, possibly making riverine ecosystems even more vulnerable to environmental changes than already known (Vörösmarty et al., 2010; Grill et al., 2015; Altermatt, 2013).

### Connectivity impacts spatial kin structure and dispersal evolution

The transition from spatially implicit (fully connected, i.e., following classical Levin’s type dynamics) to spatially explicit connectivity patterns (nearest-neigbour, circular; no variation in connectivity) has clear consequences for the cost-benefit balance underlying dispersal evolution (Fig. 1). While the costs of dispersal do not change, the benefits of dispersal do. More specifically, the probability of encountering related individuals (kin) after dispersal is altered (Fig. 2): trivially, a disperser’s chance of encountering related individuals from its patch of origin increases with deceasing connectivity, since the dispersers from a given patch of origin are dispersed to fewer target patches. The important effect of kin competition for the evolution of dispersal is well known since the seminal work of Hamilton and May (1977).

As the selective effect of connectivity on dispersal is due to the spatial correlation of kin structure, a possible adaptation to escape from such a situation would be long-distance dispersal. In our model, dispersal distance or the shape of the dispersal kernel cannot evolve (but see, e.g., Fronhofer et al., 2014, 2015). Critically, this would reduce genetic structure (*F_ST_*), increase rescue effects and, therefore, occupancy, which would reduce the occurrence of classical metapopulation dynamics.

### Dendritic topology selects against dispersal

The selective effect of network topology has recently been demonstrated by Henriques-Silva et al. (2015) for density-dependent dispersal. As expected, this also holds for density-independent dispersal strategies (Fig. 4). The mechanism behind the evolution of reduced dispersal in dendritic networks is linked to emerging and predictable heterogeneities in population densities, and, therefore, fitness (Fig. 5). Less connected patches characteristically have lower population densities and are typically connected to patches with higher population densities due to asymmetries in the number of dispersers linked to variation in connectivity as described in the results. Importantly, these density patterns and the resulting distribution of fitness in a network are spatio-temporally invariable which selects against dispersal.

Low dispersal abilities and behavioral mechanisms preventing dispersal are well-known empirically for many riverine organisms that typically live in dendritic, spatially structured populations. For example, there is a strong tendency of aquatic macroinvertebrates to escape passive drift (e.g., Elliott, 2003), and many aquatic macroinvertebrates have flight strategies in their adult stage to compensate larval downstream drift and thus reduce effective dispersal. The relatively low dispersal ability of riverine organisms is also reflected in commonly high genetic differentiation among local populations (e.g., Westram et al., 2013).

### Classical metapopulation dynamics emerge in dendritic networks

Both, reduced connectivity and dendritic topology lead to spatio-temporally correlated variation in fitness expectations, which strongly selects against dispersal. Lower ES dispersal rates lead to reduced rescue effects, which, together with some environmental stochasticity, leads to the emergence of patch extinctions. As a consequence, occupancies and turnover are intermediate, and genetic differentiation (*F_ST_*) is increased. Additionally, metapopulation persistence decreases as predicted by Vuilleumier and Possingham (2006) and discussed in detail by Gilarranz and Bascompte (2012).

Altogether, dendritic networks lead to an increased probability of observing classical metapopulation dynamics (Levins, 1969; Hanski and Gaggiotti, 2004) which are thought to be characterized by intermediate occupancies, some turnover and a more or less clear genetic differentiation between local populations (Fronhofer et al., 2012). In a theoretical study, assuming global or nearest-neigbour dispersal, as a majority of metapopulation studies do, Fronhofer et al. (2012) showed that such classical metapopulation dynamics only rarely occur in parameter space, which is in good accordance with the empirical scarcity of such classical metapopulations (among others, Baguette, 2004; Driscoll, 2007, 2008; Driscoll et al., 2010). We here report that the occurrence of classical metapopulation dynamics may be tightly linked to the underlying landscape topology, with dendritic spatially structured populations being more likely to exhibit classical metapopulation dynamics. Importantly, the exact values assumed for occupancy, turnover and genetic structure are not relevant for these conclusions as the transitions are very steep as depicted in Fig. 1.

### Classical metapopulations can likely be found in riverine systems

Among dendritic systems, riverine systems are also characterized by directional flow of water and a typical hierarchical distribution of carrying capacities (Rodriguez-Iturbe and Rinaldo, 1997; Carrara et al., 2014). Our results (Figs. 4 and 6) clearly show that adding these two features reinforces the patterns described above. Therefore, our model predicts that species living in rivers are especially likely to show classical metapopulation dynamics (Fig. 6). As for connectivity and topology, the effect of variation in carrying capacities and directional flow can be explained by an eco-evolutionary feedback linked to the evolution of dispersal: variation in carrying capacities typically selects against dispersal (Poethke et al., 2011) and the directionality of water flow leads to an increased probability of dispersal towards more connected and denser patches, which should also lead to lower ES dispersal rates.

Our theoretical prediction is in good agreement with recently reported empirical results suggesting the occurrence of metapopulation dynamics in riverine ecosystems, in a wide range of taxa, from plants, to invertebrates and vertebrates (Perkin and Gido, 2012; Göthe et al., 2012; Kuglerová et al., 2015). Evidently, dendritic connectivity is not limited to rivers. Montane terrestrial systems characterized by valleys or other habitats that are typically dendritic, like hedgerows, caves or transportation networks (see Grant et al., 2007), can exhibit similar dynamics.

### Conclusions

We analysed the evolutionary dynamics of dispersal in dendritic and other types of networks, and related these effects to the emergence of classical metapopulation dynamics. Our results illustrate ecoevolutionary feedbacks, in which landscape topology changes the evolutionarily stable dispersal strategy, which in turn feeds back on landscape level metrics link occupancy, turnover and genetic structure. Characteristically, dendritic connectivities are predestined for the emergence of classical metapopulation dynamics.

Our work has potentially important consequences for conservation: First, classical metapopulation dynamics are likely to occur in dendritic landscapes. Second, these specific dynamics are typically linked to an increased probability of extinction. This implies that populations living in dendritic landscapes, such as rivers, may be in specific need of intense and adequate conservation measures. Such measures should especially take into account anthropogenic interventions affecting connectivity and fragmentation, such as dam-and channel building (Grill et al., 2015; Grant et al., 2012). Our work indicates that riverine ecosystems, and populations in branching networks in general, may not only be threatened by changes in local conditions (Vörösmarty et al., 2010), such as habitat modifications, but also, and maybe especially, by altered large-scale landscape attributes and the resulting eco-evolutionary feedbacks.

## Acknowledgements

E.A.F. and F.A. thank Eawag and the Swiss National Science Foundation (grant no. PP00P3_150698 to FA) for funding.

## Appendix

**Figure A1:**
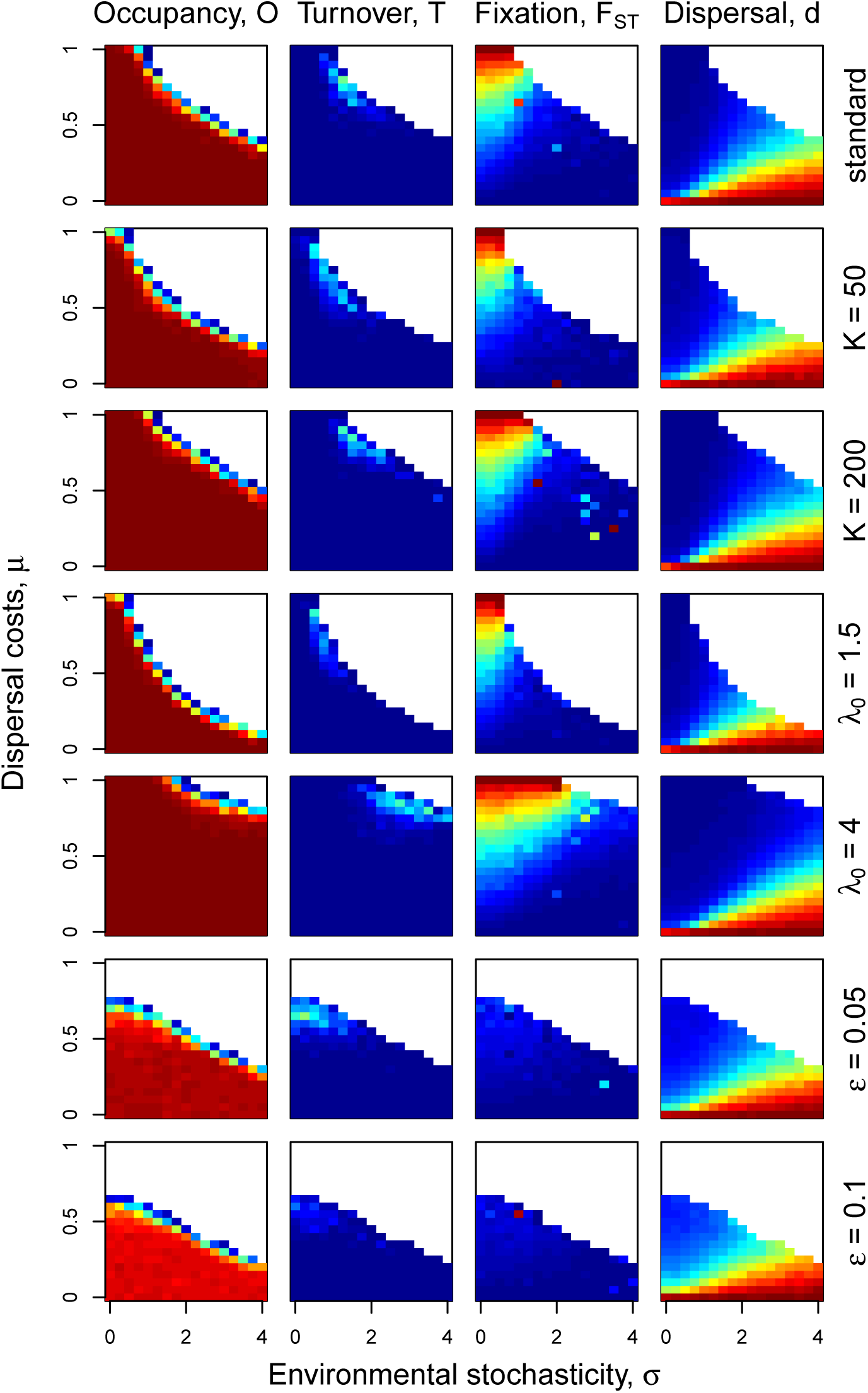
Sensitivity analysis: fully connected network. The standard parameter values were chosen to be: *K* = 100, *ϵ* = 0, *λ*_0_ = 2 (upper row). All other rows show the effect of changing one of these values while keeping the other constant. Blue colors indicate low values, red colors indicate high values, respectively. For occupancy (*O*) and fixation (*F_ST_*) values go from 0 to 1. For the ES dispersal rate (*d*) values lie between 0 and 0.98. Turnover values (*T*) are distributed between 0 and 0.12. The color coding is identical throughout the Appendix.

**Figure A2:**
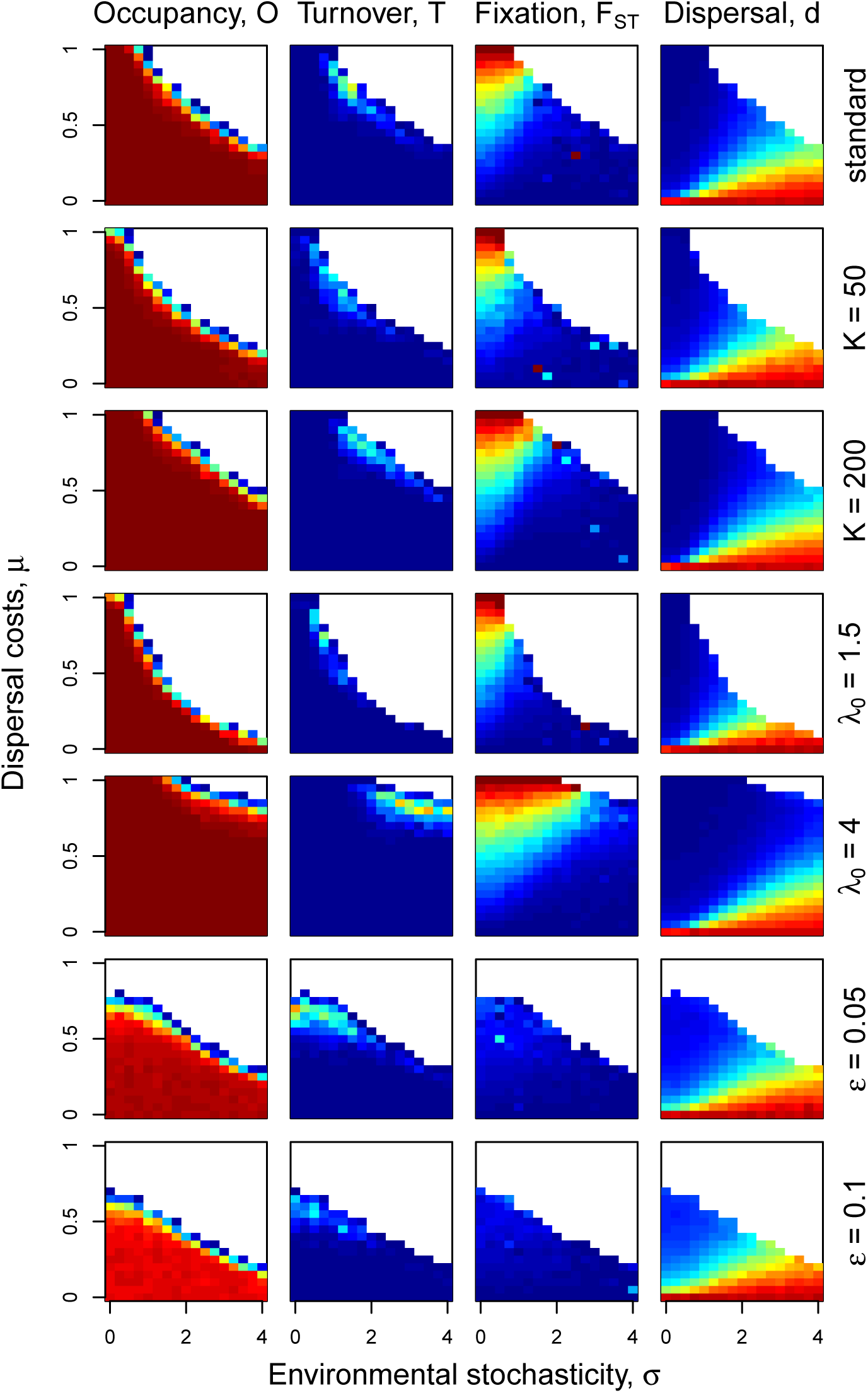
Sensitivity analysis: NN8. The standard parameter values were chosen to be: *K* = 100, *ϵ* = 0, *λ*_0_ = 2 (upper row). All other rows show the effect of changing one of these values while keeping the other constant. Blue colors indicate low values, red colors indicate high values, respectively. For occupancy (*O*) and fixation (*F_ST_*) values go from 0 to 1. For the ES dispersal rate (*d*) values lie between 0 and 0.98. Turnover values (*T*) are distributed between 0 and 0.12. The color coding is identical throughout the Appendix.

**Figure A3:**
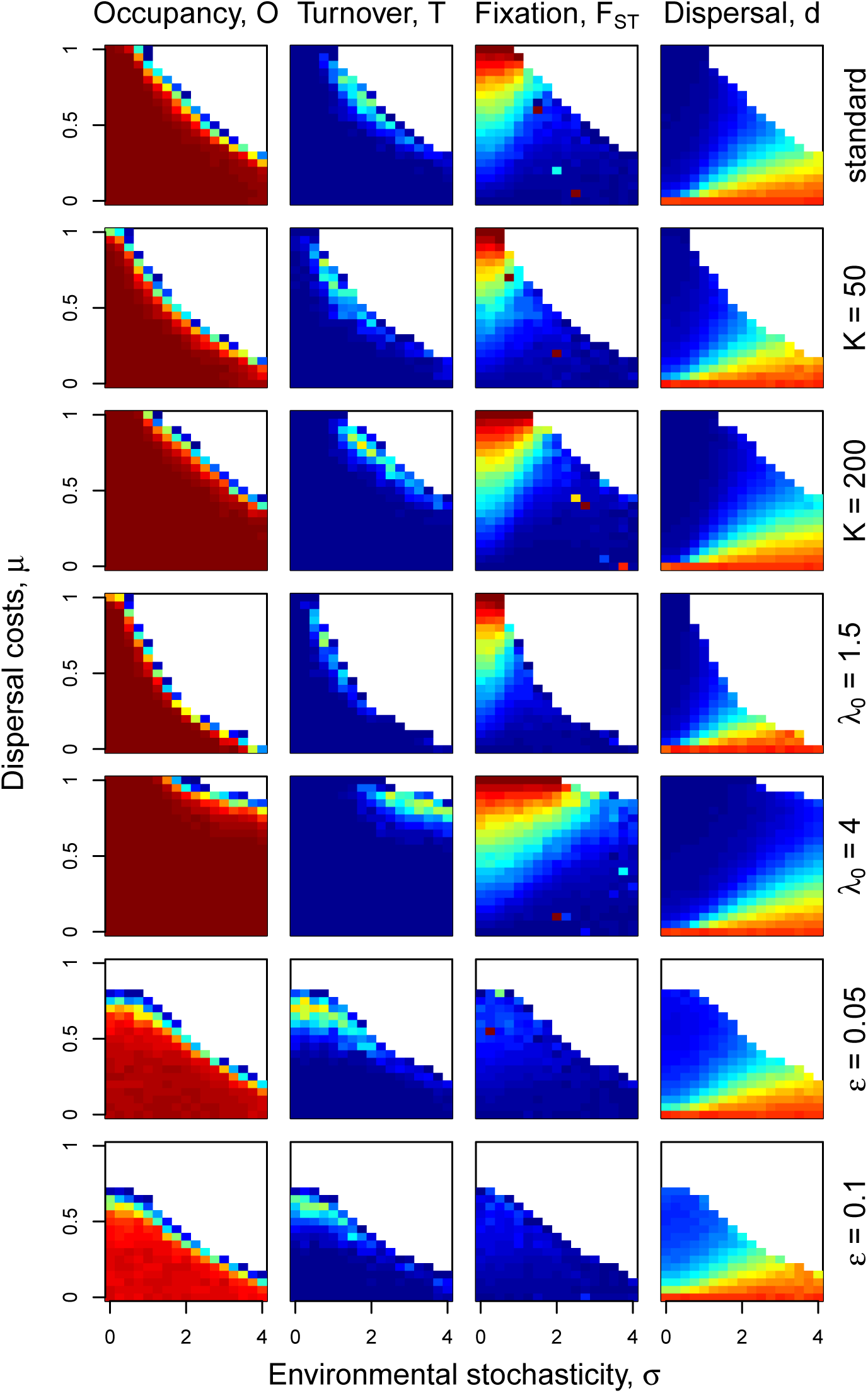
Sensitivity analysis: NN4. The standard parameter values were chosen to be: *K* = 100, *ϵ* = 0, *λ*_0_ = 2 (upper row). All other rows show the effect of changing one of these values while keeping the other constant. Blue colors indicate low values, red colors indicate high values, respectively. For occupancy (*O*) and fixation (*F_ST_*) values go from 0 to 1. For the ES dispersal rate (*d*) values lie between 0 and 0.98. Turnover values (*T*) are distributed between 0 and 0.12. The color coding is identical throughout the Appendix.

**Figure A4:**
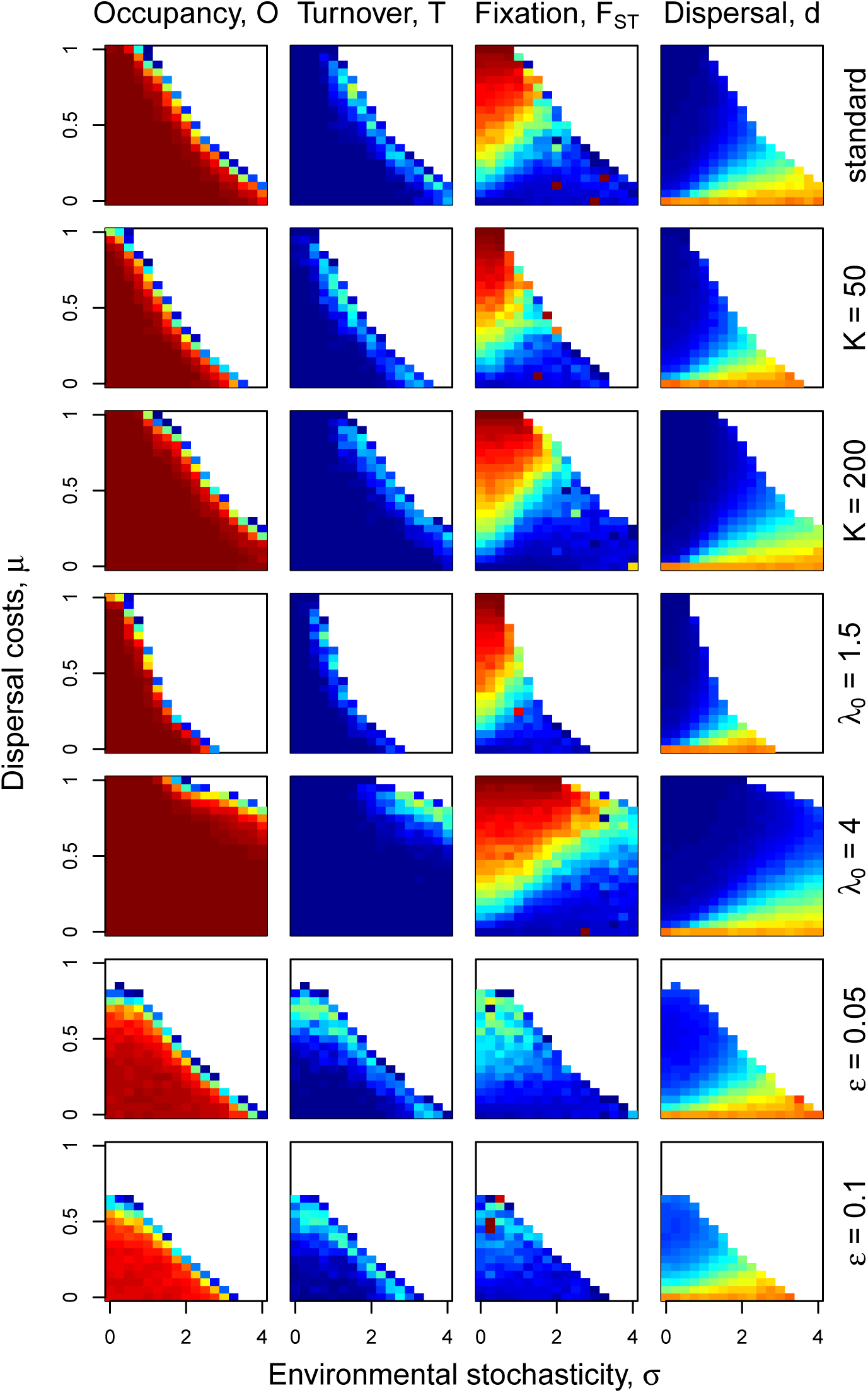
Sensitivity analysis: Circular. The standard parameter values were chosen to be: *K* = 100, *ϵ* = 0, *λ*_0_ = 2 (upper row). All other rows show the effect of changing one of these values while keeping the other constant. Blue colors indicate low values, red colors indicate high values, respectively. For occupancy (*O*) and fixation (*F_ST_*) values go from 0 to 1. For the ES dispersal rate (*d*) values lie between 0 and 0.98. Turnover values (*T*) are distributed between 0 and 0.12. The color coding is identical throughout the Appendix.

**Figure A5:**
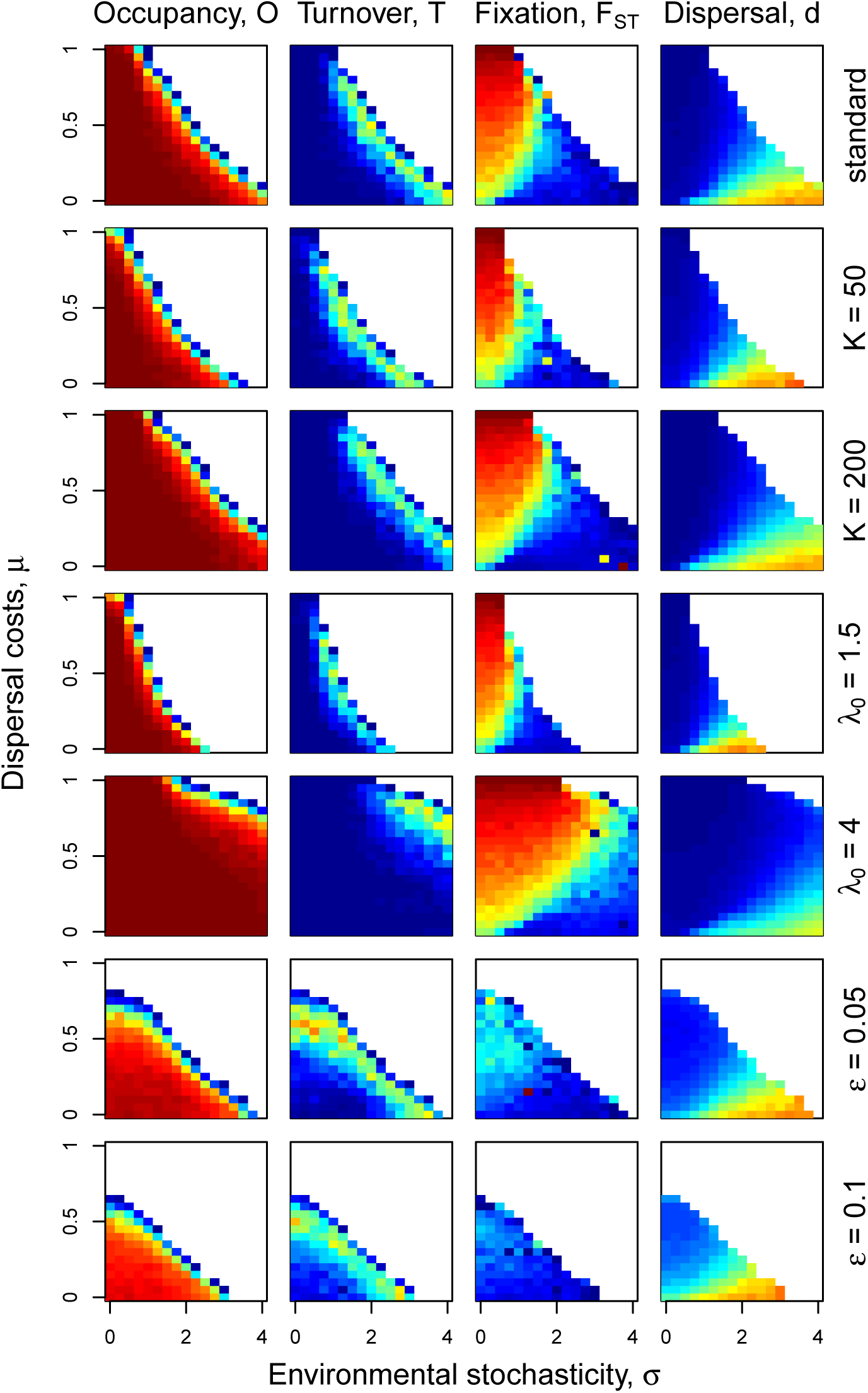
Sensitivity analysis: Dendritic bifurcated. The standard parameter values were chosen to be: *K* = 100, *ϵ* = 0, *λ*_0_ = 2 (upper row). All other rows show the effect of changing one of these values while keeping the other constant. Blue colors indicate low values, red colors indicate high values, respectively. For occupancy (*O*) and fixation (*F_ST_*) values go from 0 to 1. For the ES dispersal rate (*d*) values lie between 0 and 0.98. Turnover values (*T*) are distributed between 0 and 0.12. The color coding is identical throughout the Appendix.

**Figure A6:**
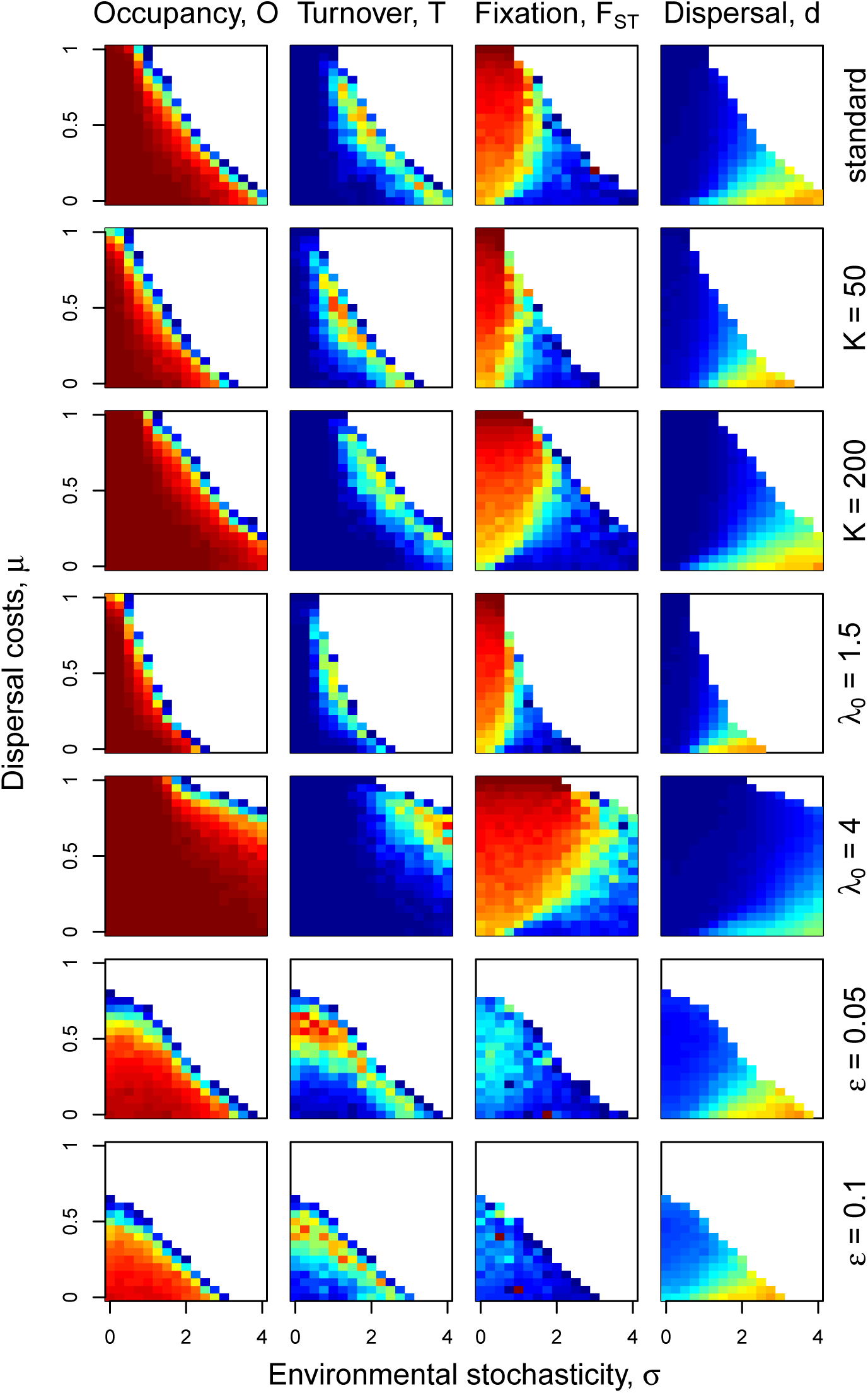
Sensitivity analysis: OCN, realization 1 (as in main text). The standard parameter values were chosen to be: *K* = 100, *ϵ* = 0, *λ*_0_ = 2 (upper row). All other rows show the effect of changing one of these values while keeping the other constant. Blue colors indicate low values, red colors indicate high values, respectively. For occupancy (*O*) and fixation (*F_ST_*) values go from 0 to 1. For the ES dispersal rate (*d*) values lie between 0 and 0.98. Turnover values (*T*) are distributed between 0 and 0.12. The color coding is identical throughout the Appendix.

**Figure A7:**
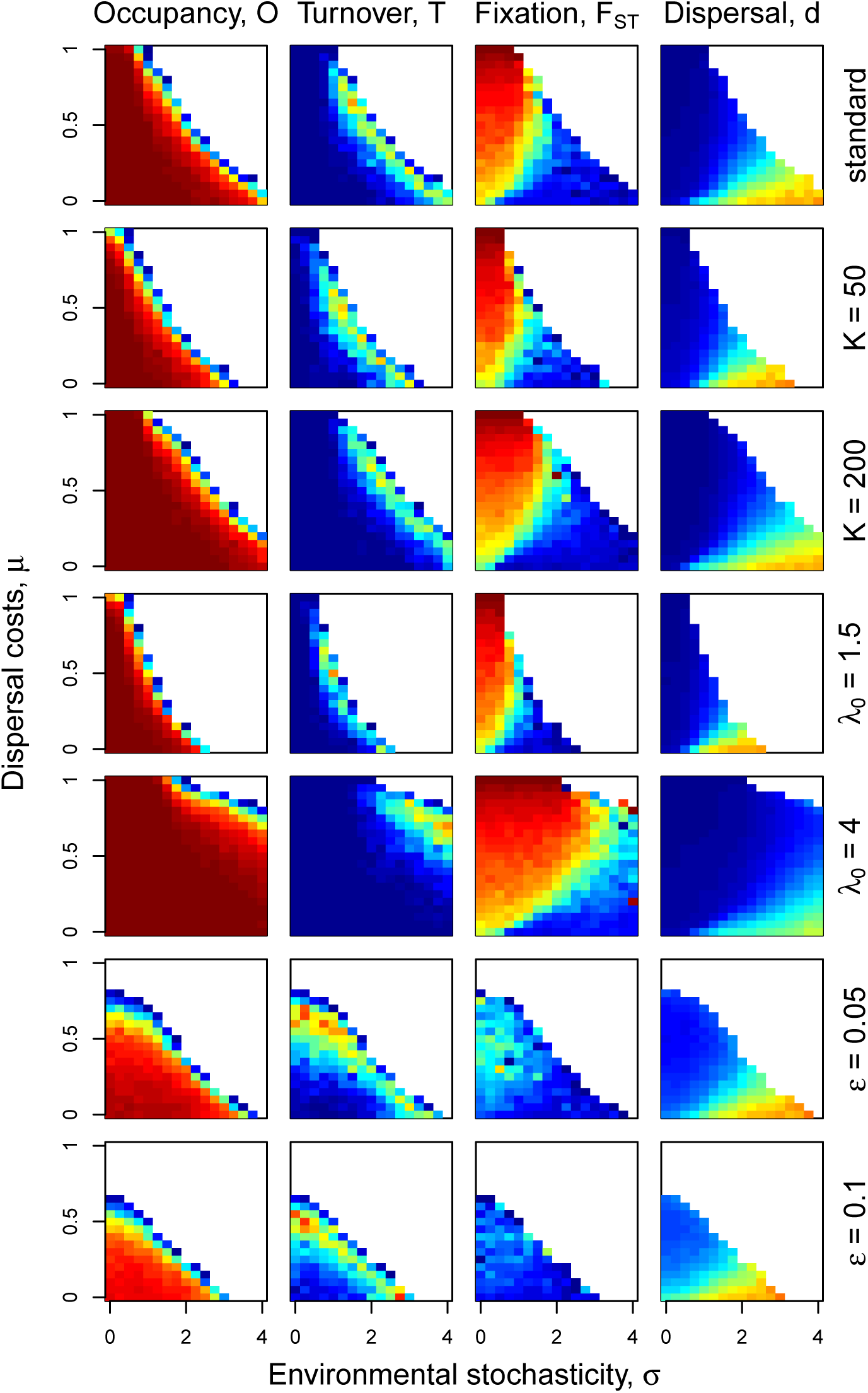
Sensitivity analysis: OCN, realization 2. The standard parameter values were chosen to be: *K* = 100, *ϵ* = 0, *λ*_0_ = 2 (upper row). All other rows show the effect of changing one of these values while keeping the other constant. Blue colors indicate low values, red colors indicate high values, respectively. For occupancy (*O*) and fixation (*F_ST_*) values go from 0 to 1. For the ES dispersal rate (*d*) values lie between 0 and 0.98. Turnover values (*T*) are distributed between 0 and 0.12. The color coding is identical throughout the Appendix.

**Figure A8:**
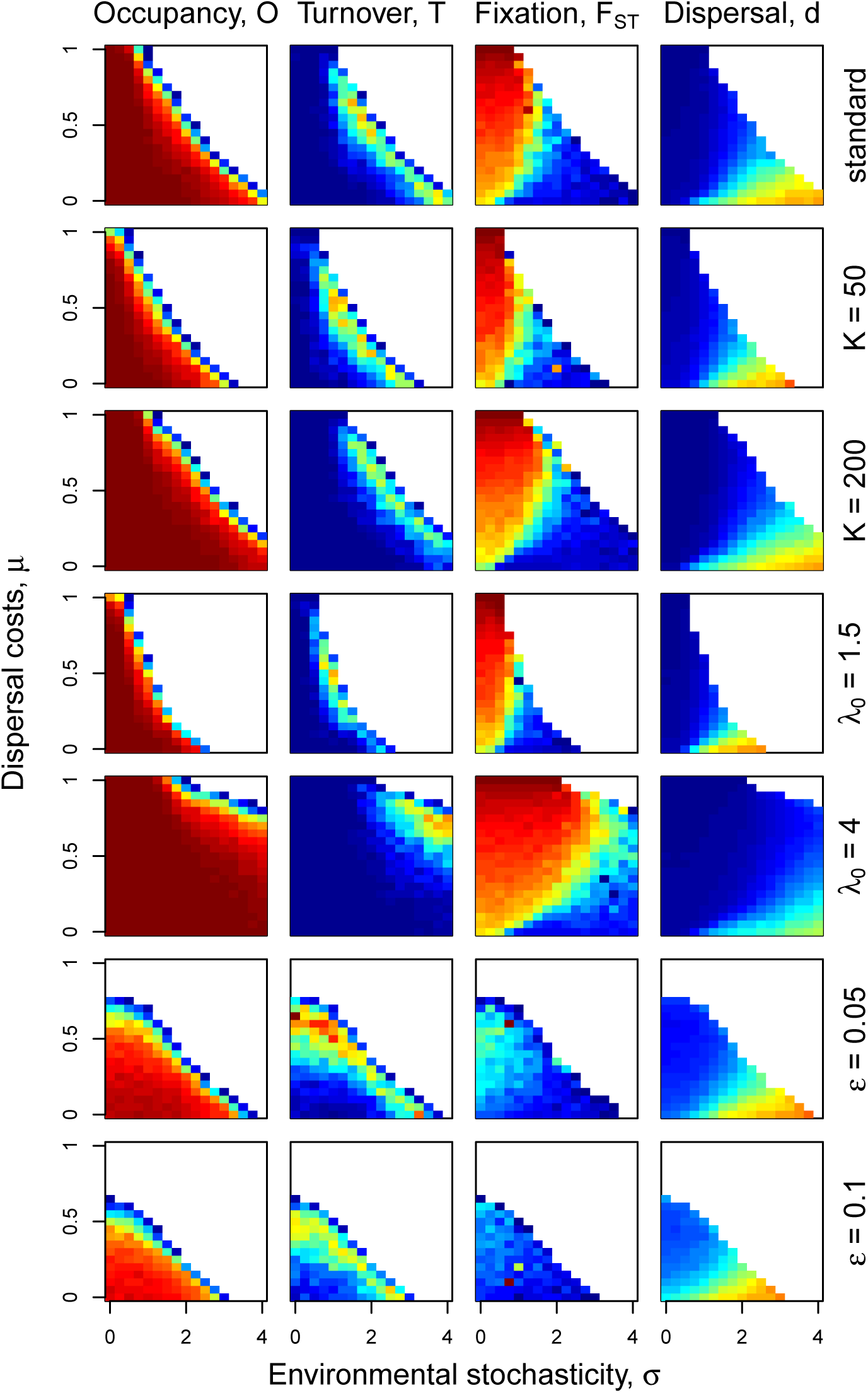
Sensitivity analysis: OCN, realization 3. The standard parameter values were chosen to be: *K* = 100, *ϵ* = 0, *λ*_0_ = 2 (upper row). All other rows show the effect of changing one of these values while keeping the other constant. Blue colors indicate low values, red colors indicate high values, respectively. For occupancy (*O*) and fixation (*F_ST_*) values go from 0 to 1. For the ES dispersal rate (*d*) values lie between 0 and 0.98. Turnover values (*T*) are distributed between 0 and 0.12. The color coding is identical throughout the Appendix.

**Figure A9:**
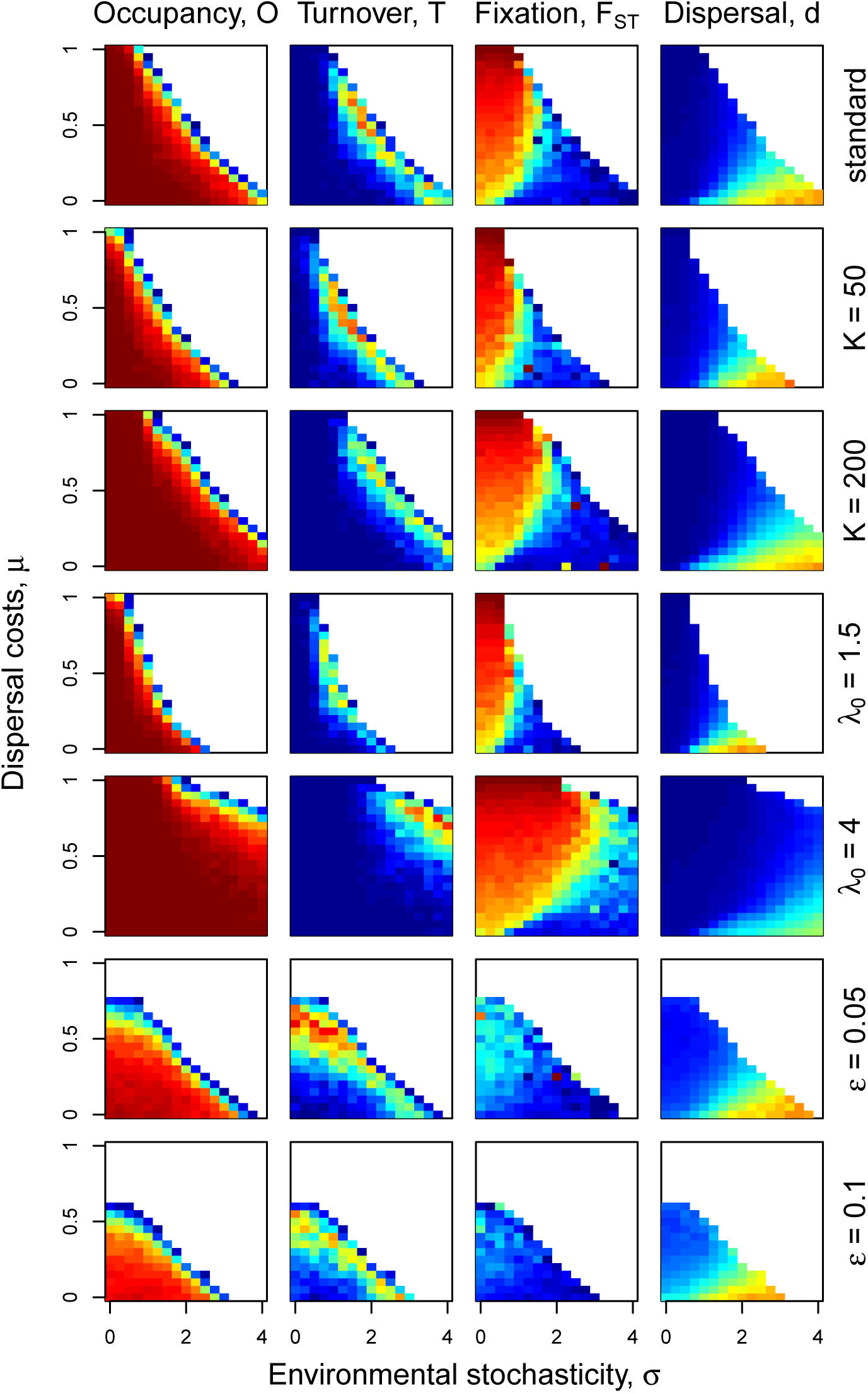
Sensitivity analysis: OCN, realization 4. The standard parameter values were chosen to be: *K* = 100, *ϵ* = 0, *λ*_0_ = 2 (upper row). All other rows show the effect of changing one of these values while keeping the other constant. Blue colors indicate low values, red colors indicate high values, respectively. For occupancy (*O*) and fixation (*F_ST_*) values go from 0 to 1. For the ES dispersal rate (*d*) values lie between 0 and 0.98. Turnover values (*T*) are distributed between 0 and 0.12. The color coding is identical throughout the Appendix.

**Figure A10:**
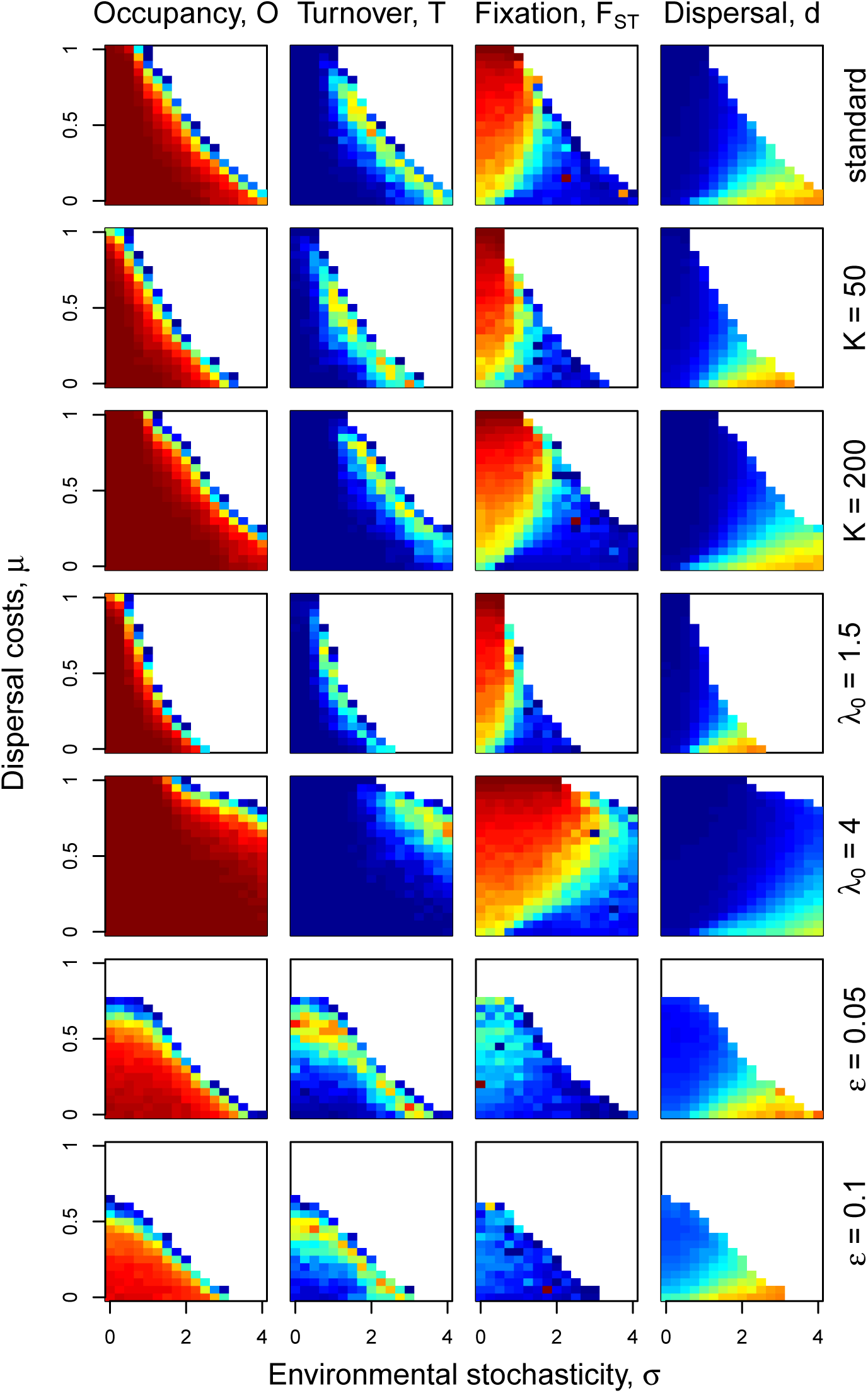
Sensitivity analysis: OCN, realization 5. The standard parameter values were chosen to be: *K* = 100, *ϵ* = 0, *λ*_0_ = 2 (upper row). All other rows show the effect of changing one of these values while keeping the other constant. Blue colors indicate low values, red colors indicate high values, respectively. For occupancy (*O*) and fixation (*F_ST_*) values go from 0 to 1. For the ES dispersal rate (*d*) values lie between 0 and 0.98. Turnover values (*T*) are distributed between 0 and 0.12. The color coding is identical throughout the Appendix.

